# Altered enhancer-promoter interaction leads to *MNX1* expression in pediatric acute myeloid leukemia with t(7;12)(q36;p13)

**DOI:** 10.1101/2023.09.13.557546

**Authors:** Dieter Weichenhan, Anna Riedel, Etienne Sollier, Umut H. Toprak, Joschka Hey, Kersten Breuer, Justyna A. Wierzbinska, Aurore Touzart, Pavlo Lutsik, Marion Bähr, Anders Östlund, Tina Nilsson, Susanna Jacobsson, Marcel Edler, Ahmed Waraky, Yvonne Lisa Behrens, Gudrun Göhring, Brigitte Schlegelberger, Clemens Steinek, Hartmann Harz, Heinrich Leonhardt, Anna Dolnik, Dirk Reinhard, Lars Bullinger, Lars Palmqvist, Daniel B. Lipka, Christoph Plass

## Abstract

Acute myeloid leukemia (AML) with the t(7;12)(q36;p13) translocation occurs only in very young children and has a poor clinical outcome. The expected oncofusion between breakpoint partners (*MNX1* and *ETV6*) has only been reported in a subset of cases. However, a universal feature is the strong transcript and protein expression of MNX1, a homeobox transcription factor that is normally not expressed in hematopoietic cells. Here, we map the translocation breakpoints on chromosomes 7 and 12 in affected patients to a region proximal to *MNX1* and either introns 1 or 2 of *ETV6*. The frequency of *MNX1* overexpression in pediatric AML (n=1556, own and published data) is 2.4% and occurs predominantly in t(7;12)(q36;p13) AML. Chromatin interaction assays in a t(7;12)(q36;p13) iPSC cell line model unravel an enhancer-hijacking event that explains *MNX1* overexpression in hematopoietic cells. Our data suggest that enhancer-hijacking is a more common and overlooked mechanism for structural rearrangement-mediated gene activation in AML.

**Key points:**

- Expression analysis of over 1500 pediatric AML samples demonstrates *MNX1* expression as a universal feature of t(7;12)(q36;p13) AML as well as in rare cases without t(7;12)(q36;p13)
- *MNX1* is activated by an enhancer-hijacking event in t(7;12)(q36;p13) AML and not, as previously postulated, by the creation of a *MNX1*::*ETV6* oncofusion gene.

## Introduction

Acute myeloid leukemia (AML) has been successfully investigated in the past using cytogenetic analysis. This led to the discovery of numerous recurrent chromosomal translocations (e.g. t(8;21)(q22; q22) or t(15;17)(q22;q12)) generating oncofusion proteins (e.g. RUNX1::RUNX1T1 or PML::RARA, respectively) that drive leukemogenesis. For many years, these translocations have served as diagnostic and prognostic markers, and affected patients can now be treated with specific targeted therapies (e.g. retinoic acid and arsenic trioxide in t(15;17) cases).^1^ In 1998, two publications reported the translocation t(7;12)(q36;p13) in AML of infants ^2,3^ occurring predominantly in children <18 months and not in adult AML. A recent meta-analysis of the Nordic Society for Pediatric Hematology and Oncology (NOPHO-AML) determined that t(7;12)(q36;p13) AML constituted 4.3% of all children with AML under the age of 2 and found a 3-year event-free survival of 24% (literature-based data) and 43% (NOPHO-AML data).^4^ Cytogenetically, t(7;12)(q36;p13) AML is often associated with the occurrence of trisomy 19 ^4,5^, but no other recurrent aberrations have been described.

Reported breakpoints in t(7;12)(q36;p13) AML have mainly been evaluated by fluorescent in situ hybridization analysis. The breakpoints on chromosome 12 (chr12) are located within intron 1 or 2 of *ETS variant transcription factor 6* or *ETV6* and proximal to M*otor neuron and pancreas homeobox 1* (*MNX1*) and within the 3’ end of *Nucleolar protein with MIF4G domain 1* (*NOM1*) on chr7.^6^ A *MNX1*::*ETV6* fusion transcript was described only in a subset of t(7;12)(q36;p13) AML cases.^4–7^ However, all AML cases with t(7;12)(q36;p13) have high expression of *MNX1* ^5^, suggesting a, yet unknown, mechanism of *MNX1* activation. Consistent with the activation of a silenced gene locus, a translocation of the *MNX1* locus from the nuclear periphery to the internal nucleus was seen, an observation that is in line with the idea that condensed and silent chromatin is located in the nuclear periphery.^6^ Also, the interactions of *ETV6* downstream elements with the *MNX1* locus have been postulated as possible mechanisms for *MNX1* activation.^6,8^ The first clue for the existence of possible aberrant promoter-enhancer interactions leading to *MNX1* activation came from our investigations of the GDM-1 AML cell line which harbors a t(6;7)(q23;q36) translocation. In GDM-1, the *MNX1* promoter interacts with an enhancer element from the *MYB* locus on chr6q23.^9^

Recently, Nilsson et al. reported the introduction of a translocation between chr7q36 and chr12p13, modeling the one found in t(7;12)(q36;p13) AML, into the human induced pluripotent stem cell (iPSC) line, ChiPSC22^WT^.^8^ The derivative line, ChiPSC22^t(7;12)^, can be differentiated into hematopoietic stem and progenitor cells (HSPC), and as such expresses *MNX1*, suggesting that hematopoietic enhancers play a role in *MNX1* activation. Enhancer-hijacking has initially been described as a mechanism for oncogene activation in AML with inv(3)/t(3;3)(q21q26) AML, where activation of *EVI1*, an isoform encoded from the *MDS and EVI1 complex locus* (*MECOM*), results from the repositioning of a *GATA2* enhancer.^10–12^ Enhancer-hijacking is also implicated in acute leukemia of ambiguous lineage in which translocated hematopoietic enhancers from different chromosomes are involved in activating *BCL11B*.^*13*^

Here we provide a detailed description of the molecular alterations found in six t(7;12)(q36;p13) AML patients and dissect the molecular mechanism leading to *MNX1* activation through the use of CRISPR-engineered ChiPSC22^t(7;12)^ iPSCs and HSPCs. We identify an enhancer-hijacking event activating the *MNX1* promoter via hematopoietic enhancers from the *ETV6* locus and validate this event in the iPSC/HSPC system. Our data suggest that enhancer-hijacking may be a more widespread, but so far largely unappreciated, mechanism for gene activation in AML with cytogenetic abnormalities.

## Methods

### Samples and cell lines

Pediatric leukemia samples T1, T2 and T3 (supplemental Table 1) were obtained at diagnosis after informed consent of patients’ legal guardians in accordance with the institution’s ethical review board (Essen, Hannover MHH, No. 2899). Sample T4 (supplemental Table 1) came from a pediatric AML cohort in Gothenburg, Sweden, and informed consent was obtained from the legal guardians in accordance with the local ethical review board. Human iPSC line ChiPSC22 (Cellartis/Takara Bio Europe AB) was cultivated in the feeder-free DEF-CS system (Cellartis/Takara Bio Europe) under standard conditions. Before differentiation, cells were transferred to Matrigel (CORNING) and mTeSR1 medium (StemCell Technologies Inc.) for 2-3 passages. ChiPSC22 was authenticated; this line and its derivatives were regularly tested for mycoplasma contamination using a commercial test kit (VenorGeM Classic, Minerva Biolabs).

### Differentiation to hematopoietic cells

Differentiation of ChiPSC22 was done as previously described.^8^

### Whole genome sequencing (WGS)

Genomic DNA was isolated using the Quick DNA Miniprep kit (Zymo research) according to the manufacturer’s protocol. WGS of four t(7;12)(q36;p13) AML cases T1-T4 was done in an Illumina HiSeq X Ten sequencer in one lane. Fastq files were aligned with BWA-MEM to the hs37d5_PhiX reference genome (hg19 with PhiX; all genomic coordinates referenced here refer to hg19). SNVs were called using mutect2. Because of the lack of matched germline sequences, only 52 known AML driver genes (supplemental Table 2) were screened for mutations. Structural variant (SVs) and somatic copy number alterations (SCNAs) were called using the Hartwig Medical Foundation (HMF) pipeline (https://github.com/hartwigmedical/hmftools). HMF tools were used in tumor-only mode, putative germline SVs were filtered out using a large panel of HMF-provided normals. SVs <20 kb were filtered out. Processed data of two samples of the TARGET-AML dataset (supplemental Table 1; dbGaP accession: phs000465.v22.p8) was downloaded using the globus platform.^14^

### RNA isolation, sequencing and qRT-PCR

Total RNA was isolated using the RNeasy Plus Mini kit (Qiagen). After library preparation, RNA was sequenced on NOVASEQ 6000 with 100 bp paired-end. The fastq files were processed using the nf-core ^15^ RNAseq v3.9 pipeline, with alignment performed using STAR ^16^ and quantification performed with Salmon.^17^ qRT-PCR was performed as previously described using TaqMan Universal Master Mix II with UNG (ThermoFisher Scientific, Applied Biosystems) and TaqMan Gene expression assays (ThermoFisher Scientific, Applied Biosystems, supplemental Table 3)^.8^

### Protein extraction, Western blotting and protein detection

Protein extraction, Western blotting and protein detection with antibodies (supplemental Table 4) was done as described previously. ^9^

### Expression screens

Processed RNAseq expression data were downloaded from the TARGET cohort ^37^ (both TARGET-NCI and TARGET-FHCRC; https://target-data.nci.nih.gov/Public/AML/mRNA-seq/L3/expression/BCCA/). *MNX1* is not expressed in normal hematopoietic cells, hence, in RNAseq a *MNX1* expression >0.5 transcripts per million (TPM) was considered overexpression. CEL files of the Balgobind cohort ^40^ were downloaded from GEO (GSE17855) and normalized using the affy R package (https://bioconductor.org/packages/release/bioc/html/affy.html). Log-expression values of microarray data were assumed to be normally distributed. We computed the mean and standard deviation for *MNX1* expression across all samples; those whose *MNX1* expression was higher than the mean plus three standard deviations were considered to express *MNX1* (3-sigma rule).

### Circular chromosome conformation capture (4C)

Circular chromosome conformation capture (4C) with each two million cells was done and analyzed as described ^9^ using *Hin*dIII in combination with *Dpn*II (supplemental Table 5).

### Hi-C

Hi-C libraries were prepared and analyzed as previously described ^18^ with minor modifications. One million cells were fixed at a final concentration of 1% formaldehyde in RPMI 1640 medium. Digestion was performed using *Dpn*II. Two to three Hi-C library replicates per sample were sequenced with 240 million reads per replicate. The fastq files were processed using the nf-core/hic v.2.1.0 pipeline. NeoLoopFinder was used to plot the contact maps.^19^

### Two-color FISH

Two probe sets of 73-75 oligonucleotides to target *MNX1* (chr7:156802250-156807250) and *ETV6* (chr12:12,160,800-12,165,800) were designed as described previously ^32^ (supplemental Table 6). Probes consist of 40 nucleotides genomic DNA and 30 nucleotides carrying regularly spaced ATTO594 or ATTO647N molecules (unpublished). Two-color FISH was conducted as previously described with minor adaptations.^20,21^ Undifferentiated ChiPSC22 were seeded on DEF-CS COAT-1-coated (Cellartis, Takara BioSciences) coverslips and differentiated ChiPSC22 on poly-L-lysine-coated coverslips. After washing and fixation steps, coverslips were mounted on microscopic slides with Mowiol (2.5% DABCO, pH 7.0; Carl Roth), dried for 30 min and sealed with nail polish.^22^ 3D FISH images were acquired using a Nikon TiE microscope (controller software NIS Elements, ver. 5.21.03) equipped with a Yokogawa CSU-W1 spinning-disk confocal unit (50 μm pinhole size), an Andor Borealis illumination unit, Acal BFi (Gröbenzell) laser beam combiner (405 nm/488 nm/561 nm/640 nm), Andor IXON 888 Ultra EMCCD camera, and a Nikon 100×/1.45 NA oil immersion objective. Confocal image z-stacks were acquired with a step size of 300 nm. Distances between *MNX1* and *ETV6* were measured using a semi-automatic in-house script in Fiji (v1.54f ^23^). Nuclear segmentation masks were generated by thresholding. ATTO594 and ATTO647N spots were detected in the nucleus through thresholding and filtered by their size and shape. Co-occurring signals within 25 x 25 pixels (xy) were included in the analysis. Local maxima were determined using 3D ImageJ Suite (V3.96) and the distances were calculated using 3D Euclidean distance.^24^ Only distances < 3.5 µm were considered. *P < 0.05, Wilcoxon rank sum test.

### Antibody-guided Chromatin Tagmentation (ACT-seq) and Assay for transposase-accessible chromatin by sequencing (ATAC-seq)

Genome-wide targeting and mapping of histone modifications (supplemental Table 4) and mapping of open chromatin were done by ACT-seq and ATAC-seq, respectively, as described previously.^9^ For read normalization using spiked-in yeast DNA in ACT-seq, trimmed reads were additionally aligned against the *Saccharomyces cerevisiae* R64 reference genome followed by post-alignment filtering. An ACT-seq library-specific scaling factor was obtained by calculating the multiplicative inverse of the number of filtered alignments against the yeast genome.^9^ ACT-seq peak calling was done applying MACS v.2.2.6 (https://pypi.org/project/MACS2/) with a q-value cutoff of 0.05 and default parameters using a wrapper script, callpeaks (https://www.encodeproject.org/chipseq/histone/), with settings narrowPeak and broadPeak for H3K27ac and H3K4me1, respectively. To facilitate visualization of hematopoietic-specific enhancers in the Integrative Genomics Viewer (IGV, version 2.11.7) ^25^, we generated HSPC-specific acetylated lysine 27 of histone 3 (H3K27ac) and mono-methylated lysine 4 of histone 3 (H3K4me1) bw-tracks with callpeaks using corresponding iPSC data as internal reference.

### Deletion of the enhancer region

A region of 212 kb (chr12:11798090-12011665) covering the four enhancers located closest to the breakpoint in ChiPSC22^t(7;12)^ was deleted by CRISPR/Cas9 editing as described previously ^8^ using CRISPR RNAs designed with the Alt-R Custom Cas9 crRNA Design Tool (Integrated DNA Technologies). To join the two ends by homology-directed repair, a 150 single-stranded deoxynucleotide (ssODN) was designed with 75 bases sequence homology on each side. Deletion was done in ChiPSC22^WT^ and ChiPSC22^t(7;12)^ sublines 14D7 and 24C7.^8^ The presence of the deletion on the translocated and the wildtype allele was validated by PCR using the Terra PCR Direct Polymerase Mix (Takara Bio Europe; supplemental Table 7). From line 14D7, cell line 2304B4, and from 24C7, lines 2305B10 and 2305C9 were generated.

## Results

### Whole genome sequencing of t(7;12)(q36;p13) AML

To precisely map structural rearrangements, somatic copy number alterations (SCNAs) and genetic mutations in t(7;12)(q36;p13) AML, we performed WGS of four t(7;12)(q36;p13) AML cases, T1, T2, T3 and T4 (**Figure 1A** and supplemental Table 1). We additionally used published WGS data from two samples with t(7;12)(q36;p13) from the TARGET cohort ^26^ (supplemental Table 1). The presence of t(7;12)(q36;p13) as a reciprocal balanced translocation was verified in all six samples (**Figure 1A**). The breakpoint on chr12 is located in five samples in intron 1 and in one sample in intron 2 of *ETV6*. On chr7, all breakpoints are located proximal to *MNX1*, in four cases within *NOM1*, located next to *MNX1*, and in two cases between *MNX1* and *NOM1* (**Figure 1B**). In none of these cases, an oncofusion gene between *MNX1* and *ETV6* is supported by the observed translocation breakpoints, leaving the main *MNX1* variant (RefSeq: NM_005515) unaffected by the genomic rearrangements (**Figure 1C**). Accompanying cytogenetic data (supplemental Table 1) revealed trisomy 19 in all cases, a result confirmed by SCNA analysis for all cases except for T1, for which SCNA analysis identified a trisomy 22 but no trisomy 19 (**Figure 1A**). We found mutations in common leukemia genes (supplemental Table 2), namely in *NOTCH2* (p.A2319V), *NF1* (p.P2310fs) and *PTPN11* (p.D61Y and p.E69K) (**Figure 1A**). Single nucleotide polymorphisms (SNPs) in the coding regions of *ETV6* indicated that the translocation leads in T1, T2 and T3 to monoallelic expression of the transcriptional repressor gene *ETV6* (data not shown; no RNAseq data available from T4). A previously described alternative *ETV6* exon 1B ^27^, located upstream of exon 3, is not expressed (data not shown).

**Figure 1:**
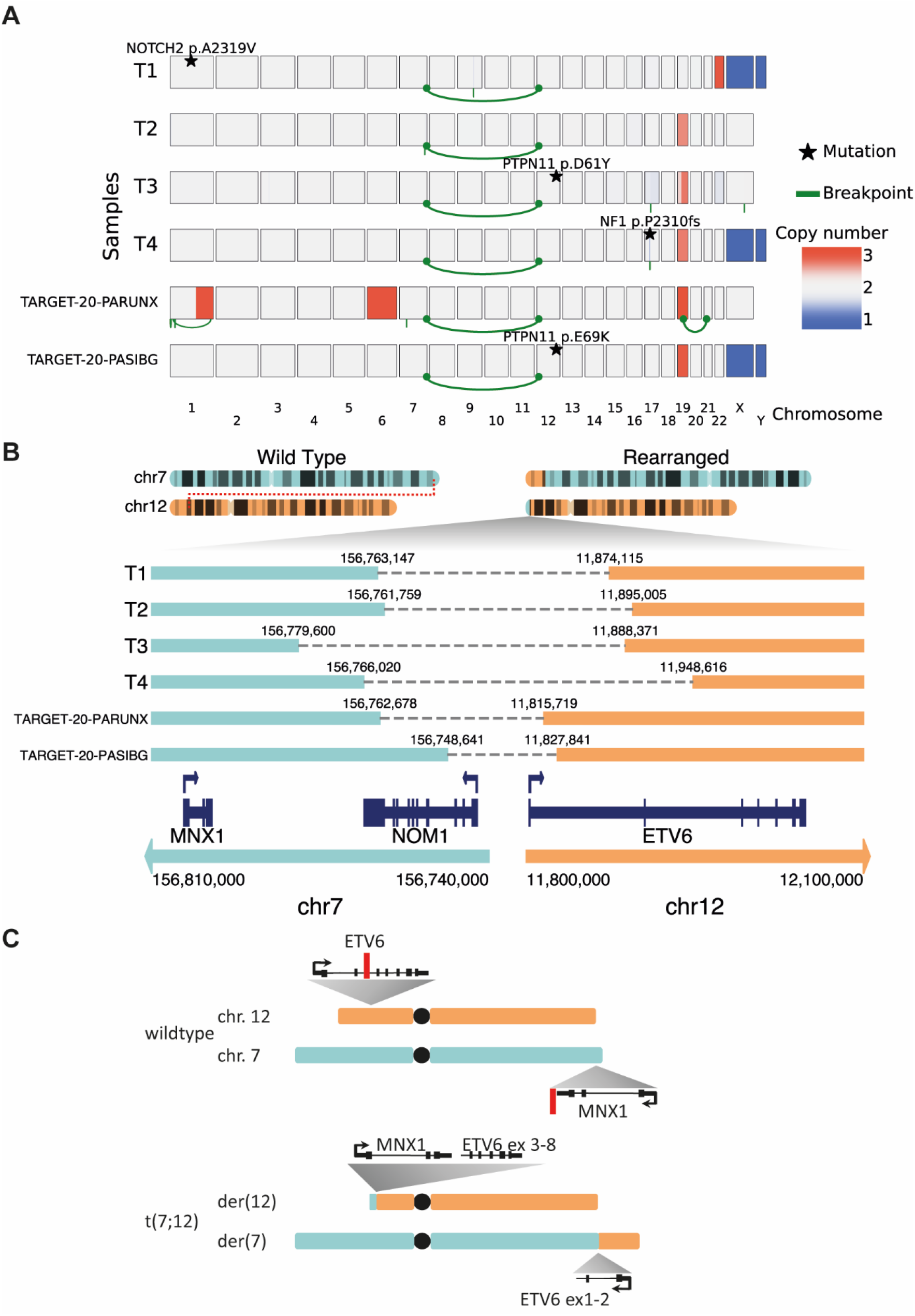
Whole genome sequence analysis of t(7;12)(q36;p13) AML. **A**, Copy numbers (blue: loss; red: gain), structural rearrangements (green bow connecting 2 chromosomes) and mutations in known AML driver genes for six t(7;12)(q36;p13) AML samples based on WGS. Samples T1, T2, T3 and T4 were profiled in this study, while TARGET-20-PARUNX and TARGET-20-PASIBG are from the TARGET-AML cohort 15. **B**, Sketch of the rearranged chromosomes 7 and 12, and zoom-in on the region around the breakpoints. **C**, Schematic overview of chr7 (turquoise), chr12 (orange) and derivative chromosomes der(12) and der(7) resulting from the reciprocal t(7;12) translocation involving *MNX1* on chr7 and *ETV6* on chr12. Red lines indicate positions of break/fusion points.

### *MNX1* is highly expressed in all t(7;12)(q36;p13) AML and is associated with a characteristic gene expression signature

Although normally not expressed in the hematopoietic lineage, *MNX*1 is highly expressed in all analyzed t(7;12)(q36;p13) AML.^5,6,28^ In line with this, AML cases T1-T4 showed high *MNX1* expression (supplemental Table 1). We additionally evaluated *MNX1* expression in two pediatric AML cohorts with available expression data: Balgobind et al. 2011 (237 samples profiled with Affymetrix array) ^29^ (**Figure 2A**) and TARGET-AML (1319 samples profiled with RNAseq) ^26^ (**Figure 2B**). *MNX1* was expressed in 7/237 (2.9%) samples of the Balgobind cohort, and in 31/1319 (2.3%; including resample for samples TARGET-20-PARUNX and TARGET-21-PASVJS) samples of the TARGET-AML cohort. All t(7;12) samples showed *MNX1* expression, but also some samples without 7q36-rearrangements. Accordingly, there might be alternative mechanisms leading to *MNX1* activation. Most t(7;12)(q36;p13) samples were diagnosed younger than 2 years old, however, most *MNX1-*overexpressing samples without t(7;12) were diagnosed at an older age (supplemental Table 1).

**Figure 2:**
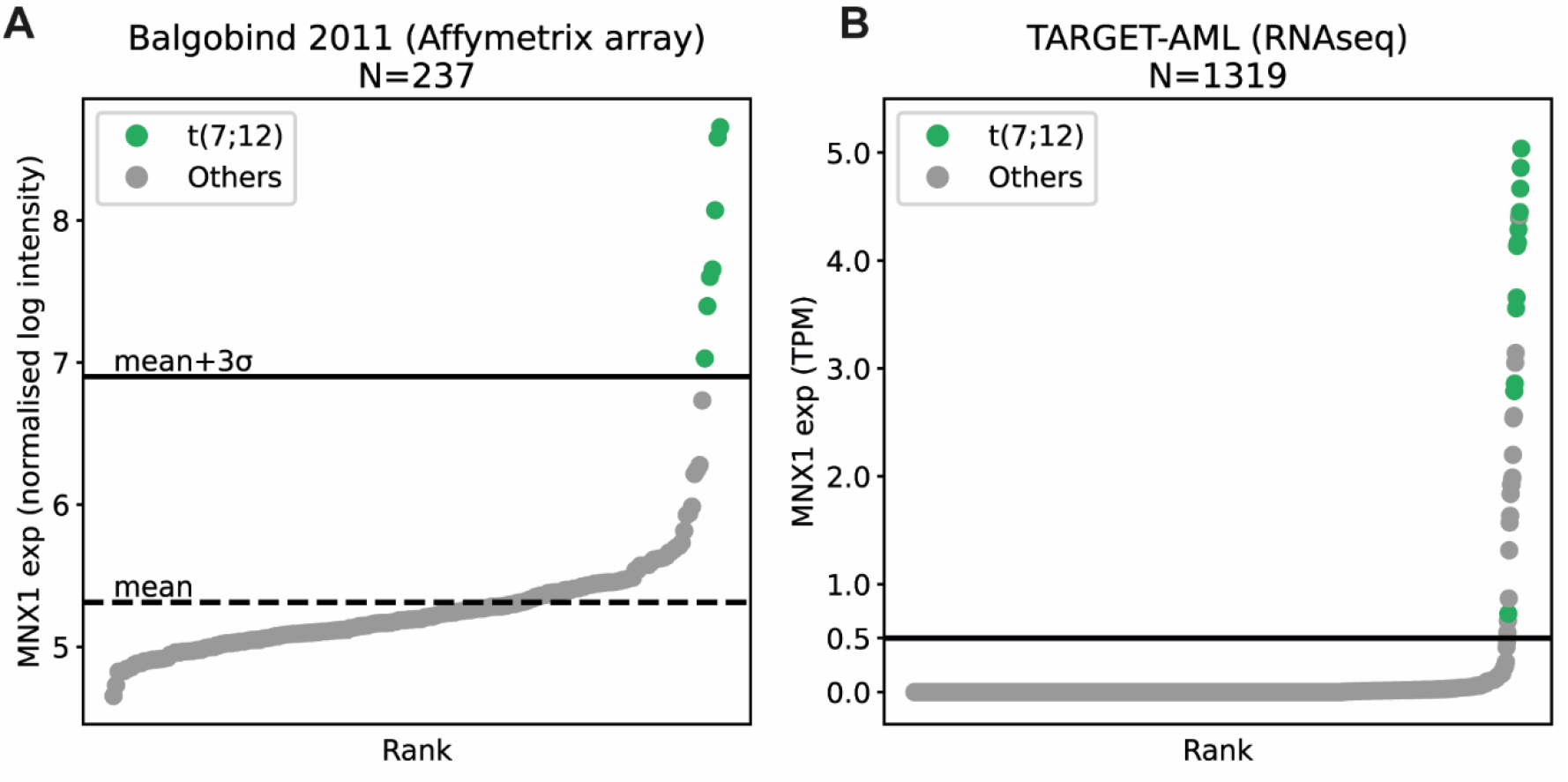
*MNX1* expression in pediatric AML with t(7;12)(q36;p13) translocation. *MNX1* expression in two different pediatric AML cohorts: **A**, Balgobind et al. 2011 ^29^, 237 samples profiled with Affymetrix arrays. The mean expression level is shown with a dashed line, and the mean plus three standard deviations is shown with a horizontal line. **B**, TARGET-AML ^26^, 1319 samples profiled with RNAseq (transcripts per million, TPM; cut-off 0.5 TPM). Samples with cytogenetically detected t(7;12)(q36;p13) translocation are shown in green and other samples in grey.

A characteristic gene expression signature for t(7;12)(q36;p13) AML as compared to other cytogenetic subgroups in pediatric AML has been described. ^29^ The majority of these genes are either consistently downregulated in t(7;12)(q36;p13) AML (e.g. *TP53BP2*) or upregulated together with *MNX1* (*EDIL3, LIN28B, BAMBI, MAF, FAM171B, AGR2, CRISP3, KRT72* and *MMP9*). We performed differential expression analysis between the t(7;12)(q36;p13) and the other cases from each the Balgobind and the TARGET-AML cohort (supplemental Table 8). Our lists of up- and downregulated genes include the genes identified by Balgobind et al. ^29^, which are indeed consistently deregulated in t(7;12)(q36;p13) across several cohorts, as well as other genes not reported before. The samples with *MNX1* expression but without genomic rearrangement close to *MXN1* did not express this typical gene signature (supplemental Figure 1). The breakpoints on chr12 are located in *ETV6* and, hence, led to a corrupted *ETV6* allele. In line with this, we found in the two cohorts a negative log-fold change of *ETV6* expression in the t(7;12)(q36;p13) samples compared to the others (data not shown).

### A t(7;12)(q36;p13) cell line model exhibits MNX1 protein expression and chromatin interactions between the *MNX1* and *ETV6* regions

Previously, Nilsson et al. engineered an iPSC line harboring a balanced translocation t(7;12)(q36;p13) (ChiPSC22^t(7;12)^) with a breakpoint in *ETV6* intron 2 and a breakpoint about 21 kb proximal to *MNX1* in the candidate common breakpoint region.^8,30^ Upon differentiation of ChiPSC22^t(7;12)^ cells to HSPCs, *MNX1* became activated, while it remained silent in the differentiated parental line ChiPSC22^WT^. We confirmed this result at the protein level by Western blot in three ChiPSC22^t(7;12)^ sublines, 14D7, 23G8 and 24C7; the MNX1 protein was only expressed in the ChiPSC22^t(7;12)^ HSPCs but neither in the ChiPSC22^t(7;12)^ iPSCs nor in ChiPSC22^WT^ iPSCs or HSPCs (**Figure 3A**). To examine if the t(7;12)(q36;p13) translocation could result in enhancers from the *ETV6* region interacting with the *MNX1* promoter, we profiled the iPSCs and HSPCs of both ChiPSC22^WT^ and ChiPSC22^t(7;12)^ lines with Hi-C. No interactions were seen between chr7 and chr12 in the ChiPSC22^WT^ (supplemental Figure 2), but we observed a new topologically associating domain (neo-TAD) around the breakpoint in both iPSC and HSPC ChiPSC22^t(7;12)^ **(Figure 3B)**. The neo-TAD extends up to chr12:12,200,000, meaning that enhancers located between the breakpoint and the end of the neo-TAD could interact with the *MNX1* promoter. To confirm these interactions, we also performed circular chromosome conformation capture (4C) using an *MNX1* viewpoint **(Figure 3B)**. We saw interactions between the *MNX1* and *ETV6* regions in HSPCs but not in iPSCs, indicating that the interactions are likely stronger in HSPCs. To strengthen the interaction data, we performed two-color FISH targeting the *MNX1* promoter and, located distal to *ETV6*, a 5 kb region which overlaps with the region that showed interaction in Hi-C. We observed a decreased 3D distance between the targeted regions in ChiPSC22t(7;12) HSPCs as compared to the corresponding iPSCs (median HSPC: 0.53 µm, median iPSCs: 1.06 µm, **Figure 3C**). The shorter distance in the HSPCs suggests a reinforced contact between an enhancer from the *ETV6* neo-TAD region and the *MNX1* promoter upon differentiation.

**Figure 3:**
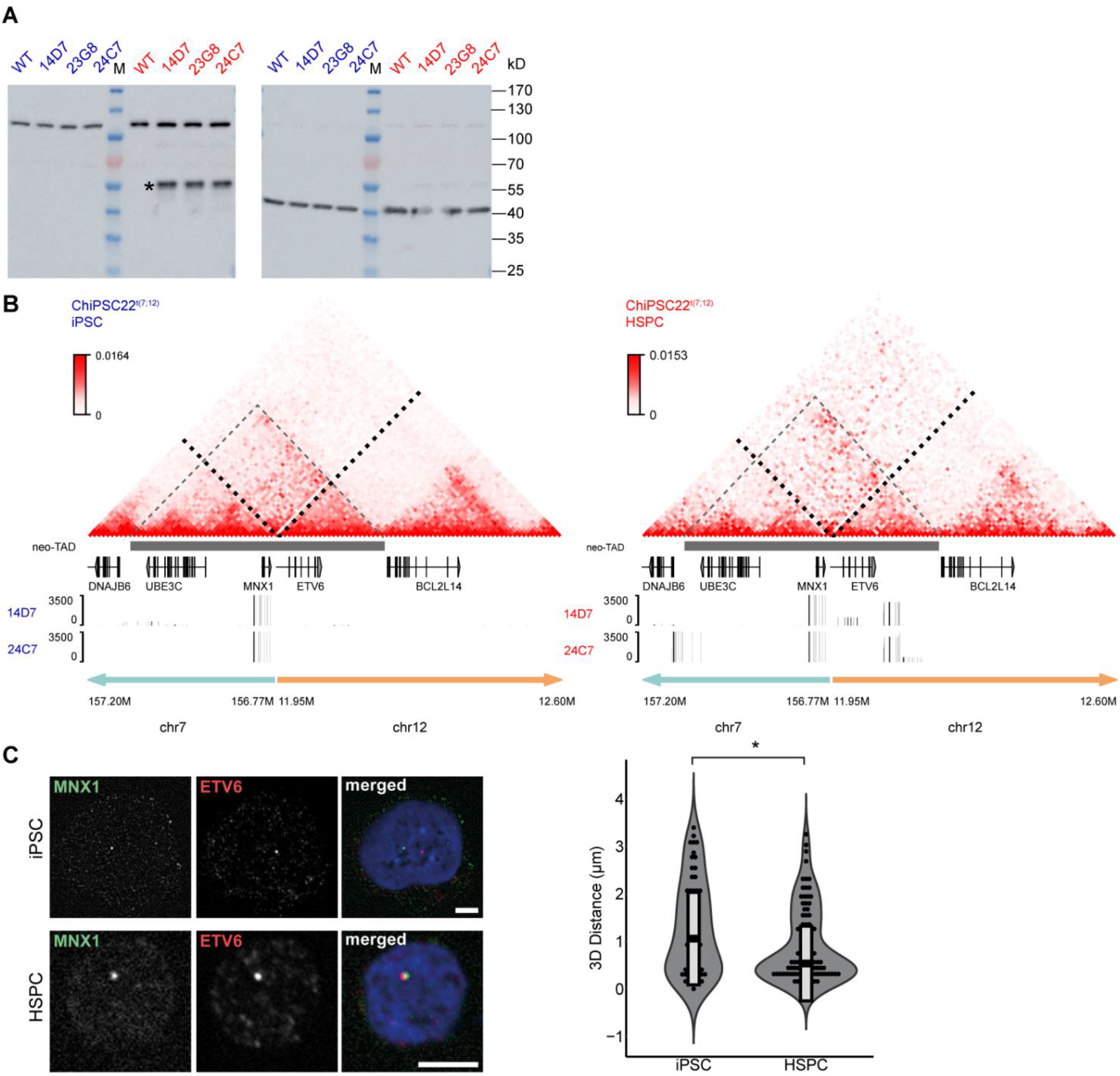
MNX1 protein expression and chromatin interaction of the *MNX1* gene with the *ETV6* region in ChiPSC22^t(7;12)^ HSPCs. **A**, Western blot with an MNX1 antibody (left) and iPSC (blue) and HSPC (red) protein extracts from ChiPSC22^WT^ and ChiPSC22^t(7;12)^ sublines 14D7, 23G8 and 24C7. The MNX1 protein (asterisk) is only detected in HSPCs of ChiPSC22^t(7;12)^ sublines 14D7, 23G8 and 24C7. The common band at about 120 kD results from an unknown protein cross-reacting with the MNX1 antibody. To demonstrate loading of equal protein amounts, the unstripped blot was re-incubated with an antibody against β-actin (right). **B**, Chromatin interactions analyzed by Hi-C seq in the genomic region flanking the translocation breakpoint in the ChiPSC22^t(7;12)^ subline 24C7, either as iPSCs (left) or HSPCs (right). The bottom tracks show 4C-data with a viewpoint about 3 kb distal to *MNX1*, for the two ChiPSC22^t(7;12)^ sublines 14D7 and 24C7. The range 0-3500 corresponds to read numbers in bins. **C**, Increased proximity between *MNX1* and *ETV6* in ChiPSC22t(7;12) derived HSPCs compared to iPSCs. Left: Representative planes from two-color FISH targeting 5 kb regions at the *MNX1* promoter and distal to *ETV6* in iPSCs and HSPCs. The iPSC image was scaled down; bars, 5 µm. Right: 3D distances between the *MNX1* and *ETV6* signals. Black horizontal lines within boxes indicate medians; box limits indicate upper and lower quartiles. iPSC: n = 64, HSPC: n = 90, across 3 independent replicates.

### Identification of hematopoietic enhancers in *ETV6* and its vicinity

Since *MNX1* expression is only seen in HSPCs but not in iPSCs, we searched for hematopoietic enhancers located in the chr12 part of the neo-TAD. Active enhancers reside in open chromatin, hence, we profiled accessible chromatin by ATAC in the two AML patient samples AML-T1 and -T2 and in HSPCs of our cell line model. We found four consistent open chromatin sites common to the patient samples and to the ChiPSC22^t(7;12)^ and ChiPSC22^WT^ cells within the *ETV6* neo-TAD, proximal to the chr12-breakpoint in the ChiPSC22^t(7;12)^ cell lines (**Figure 4**). Additionally, we mapped peaks of enhancer marks, i.e. H3K27ac and H3K4me1 in both iPSCs and HSPCs from ChiPSC22^wt^ and ChiPSC22^t(7;12)^. To facilitate the identification of hematopoietic enhancers in ChiPSC22^t(7;12)^ and ChiPSC22^WT^ HSPCs, we applied MACS2-peak calling using the corresponding iPSC data as internal reference. The four open chromatin regions were also marked by HSPC-specific H3K27ac and H3K4me1 peaks. In addition, we used public ChIPseq-datasets for the same enhancer marks (https://www.ncbi.nlm.nih.gov/geo/query/acc.cgi?acc=GSM772885; https://www.ncbi.nlm.nih.gov/geo/query/acc.cgi?acc=GSM621451) and the histone acetyltransferase P300 generated from CD34+ cells and the chronic myeloid leukemia derived cell line MOLM-1.^10^ Again the four ATAC/H3K27ac/H3K4me1 peaks were found as well in MOLM-1 and CD34+ cells (**Figure 4**). Two of these peaks, the one located closest and that one most distant to the ChiPSC22^t(7;12)^ breakpoint, coincide in addition with p300 peaks (**Figure 4**). In conclusion, we identified four strong enhancer candidates in the *ETV6* neo-TAD region, two coinciding with p300 peaks, which may drive *MNX1* activation in t(7;12)(q36;p13) AML.

**Figure 4:**
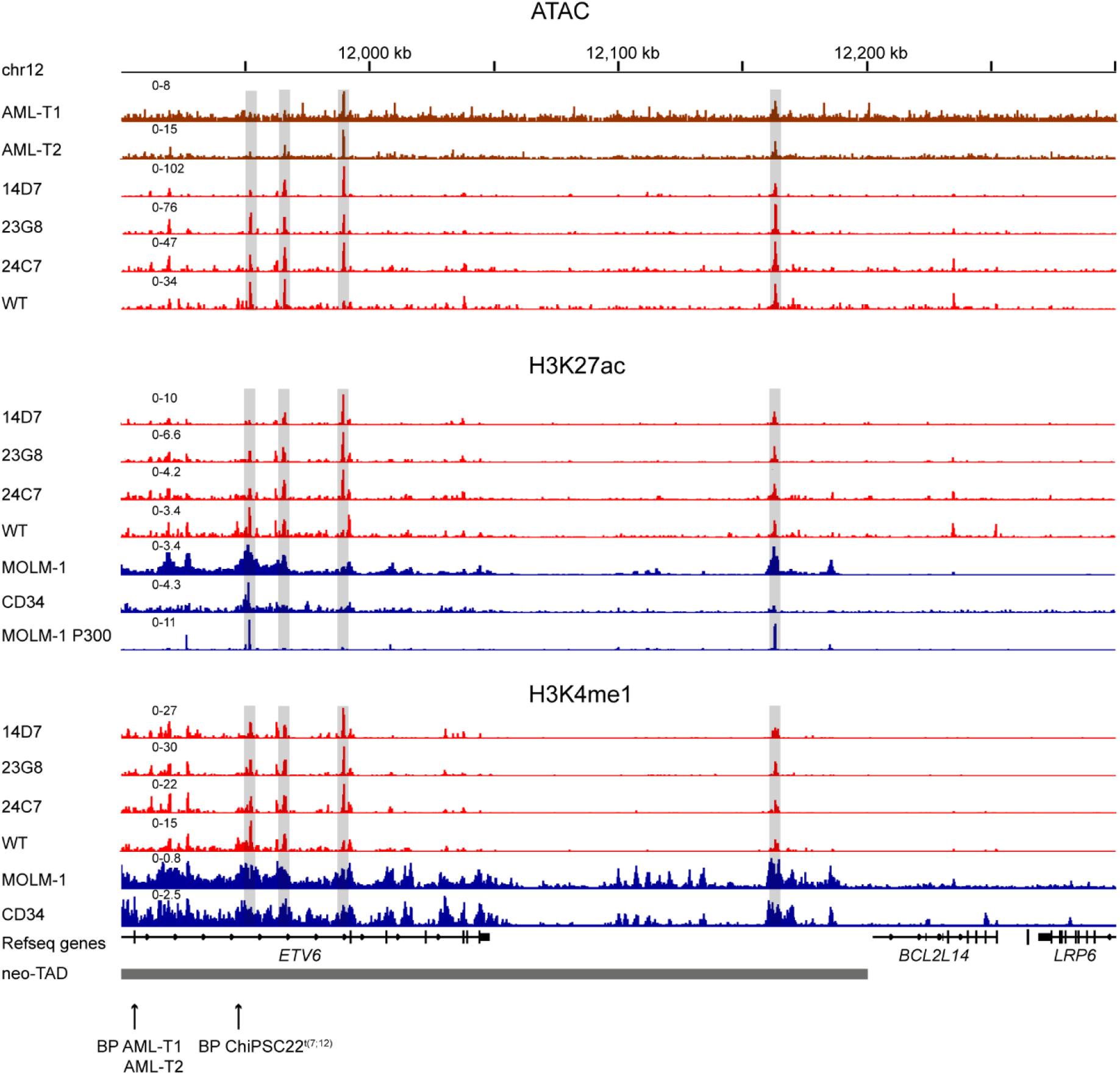
Open chromatin and enhancer mark profiles in the *ETV6* neo-TAD region of patient and cell line samples. Open chromatin profiles (ATAC) of AML patients T1 and T2 and of ChiPSC22^t(7;12)^ sublines 14D7, 23G8 and 24C7 and of ChiPSC22^WT^ HSPCs in the *ETV6* neo-TAD region. HSPC-specific H3K27ac, HSPC-specific H3K4me1 and publicly available^10^ p300, H3K4me1 and H3K27ac profiles from MOLM-1 and CD34+. Relevant common peak positions are highlighted by a gray shading. The chr12-breakpoint (BP) position in T1 and T2 and in the ChiPSC22^t(7;12)^ sublines are indicated.

### Deletion of enhancers in the *ETV6* region abrogates *MNX1* expression in ChiPSC22^t(7;12)^ HSPCs

As a further layer of experimental evidence for *de novo* promoter-enhancer interactions in t(7;12)(q36;p13) AML, we examined *MNX1* expression levels in three ChiPSC22^t(7;12)^ sublines carrying a deletion of 212 kb which removed the four enhancer candidates (ChiPSC22^t(7;12)ΔEn^, **Figure 5A**, supplemental Figure 3). In all three ChiPSC22^t(7;12)ΔEn^ lines, HSPC-specific *MNX1* expression was abrogated (**Figure 5B**). This supports the hypothesis that *MNX1* activation in ChiPSC22^t(7;12)^ is the result of interactions between the *MNX1* promoter and one or multiple enhancers located in or close to *ETV6*. We also observed a significant downregulation of genes that are upregulated together with *MNX1* in t(7;12)(q36;p13), such as *AGR2, MMP9, MAF* and *CRISP3*, suggesting that they are regulated by *MNX1*.

**Figure 5:**
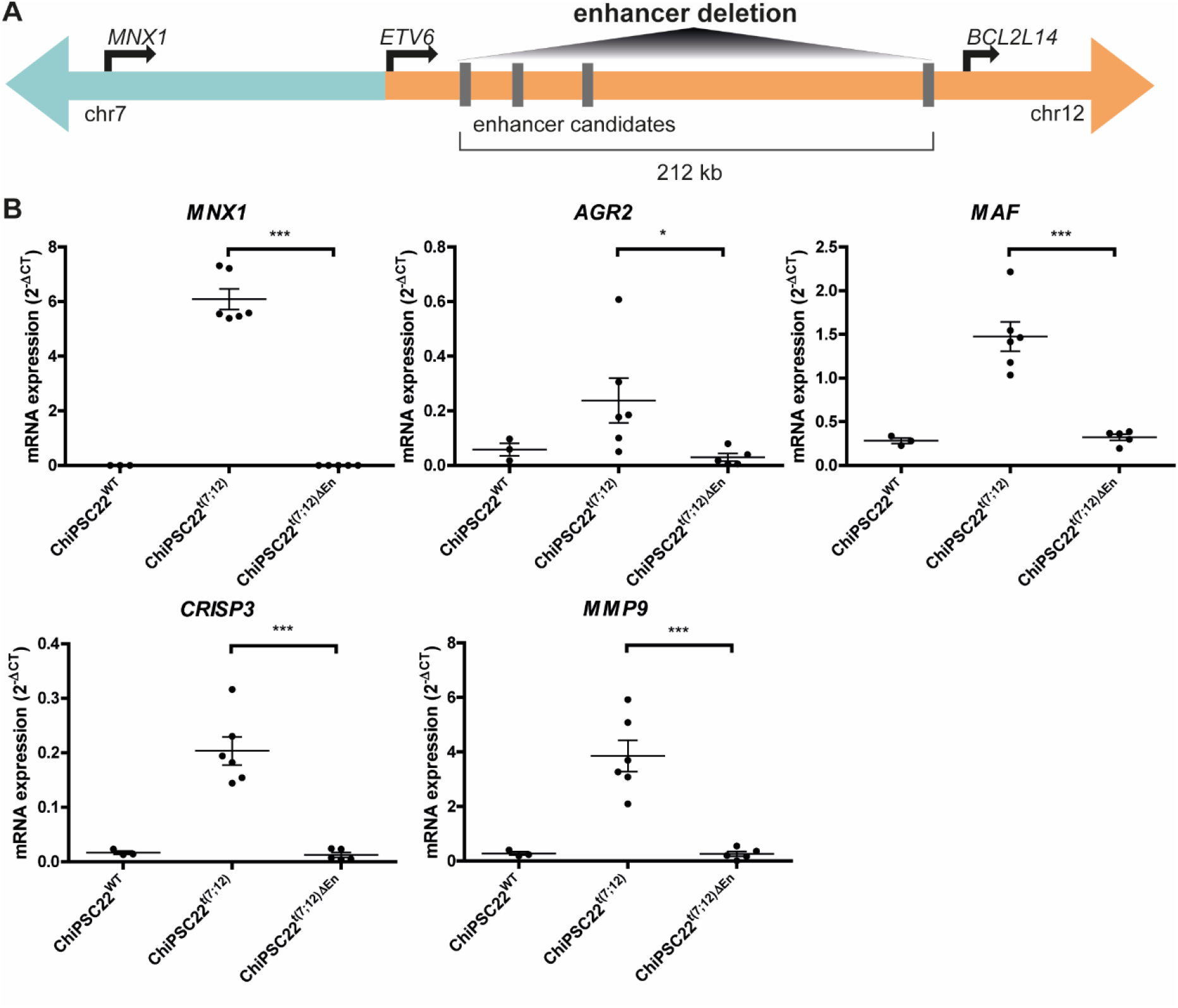
Molecular validation of enhancer-promoter interaction in ChiPSC22^t(7;12)^ upon differentiation. **A**, Scheme of experiments performed to validate the interaction between the *MNX1* promoter and enhancers distal to the breakpoint including deleting the enhancer region and spatial proximity probing. **B**, Gene expression in ChiPSC22^WT^ (n=3), ChiPSC22^t(7;12)^ (n=6, from 3 independent cell lines) and ChiPSC22^t(7;12)ΔEn^ (n=5, from 3 independent cell lines) measured via qRT-PCR and shown as 2^−ΔCt^ vs. *GUSB as* endogenous reference.

In conclusion, we provide experimental evidence for an enhancer-hijacking event rather than the creation of an oncofusion protein in pediatric AML with t(7;12)(q36;p13). Using an *in vitro* iPSC/HSPC cell system, we demonstrate that one or several enhancers in the *ETV6* region interact with *MNX1* to regulate its expression.

## Discussion

In this manuscript, we describe enhancer-hijacking and activation of *MNX1* as a novel molecular mechanism resulting from a translocation between chr7 and 12 [t(7;12)(q36;p13)] in pediatric AML. Our study shifts the focus from a putative *MNX1::ETV6* oncofusion transcript to the activation of *MNX1* as the unifying putative leukemia-driving event. Overexpression of *MNX1* is accompanied by mono-allelic inactivation of *ETV6* on the translocated chromosome, putatively resulting in haploinsufficiency. In addition, the majority of published t(7;12)(q36;p13) AML cases show trisomy 19 as a recurrent secondary event, suggesting that activation of a gene on chr19 is an additional component in the leukemogenesis of this AML subtype.^31^ Our observation has important implications for the diagnosis of this subgroup of patients, as well as novel therapeutic approaches. As shown in the expression reanalysis, all t(7;12)(q36;p13) AML demonstrate overexpression of *MNX1*. Only few AML cases without t(7;12)(q36;p13) show *MNX1* overexpression, but while t(7;12)(q36;p13) AML is diagnosed at very young age (<20 months), *MNX1* overexpression in the absence of this translocation occurs predominantly at a later age, suggesting different, yet, unexplained molecular pathways converging in *MNX1* expression. Considering that *MNX1* is not expressed in the normal hematopoietic system, quantitative *MNX1* expression analysis could be used as a diagnostic marker for this subgroup of pediatric AML.

Enhancer-hijacking events resulting in the activation of proto-oncogenes have been described also in other human malignancies including translocations resulting in the activation of oncogenes *MYC, BCL2* or *CCND1* in B-cell lymphoma ^32–34^ or rearrangements in medulloblastoma.^35^ Subtype-specific 3D genomic alterations were recently discovered in AML leading to enhancer-promoter or enhancer-silencer loops.^19^ We demonstrated that the t(7;12)(q36;p13) translocation results in a neo-TAD, where the *MNX1* promoter is able to interact with the *ETV6* region.

Initial evidence for an oncogenic role of MNX1 in leukemogenesis comes from a study by Nagel *et al*. characterizing the *MNX1*-overexpressing cell line GDM-1.^36^ Knockdown of *MNX1* led to a reduction of cell viability and cell adhesion. *In vitro* overexpression of *MNX1* in HT1080 and NIH3T3 cells leads to premature, oncogene-induced senescence mediated by the induction of p53-signalling.^37^ *In vivo*, ectopic *MNX1* expression in murine hematopoietic stem and progenitor cells resulted in strong differentiation arrest and accumulation at the megakaryocyte/erythrocyte progenitor stage.^37^ Overall, this phenotype is in line with reports on t(7;12)(q36;p13) AML blast cells which are less differentiated (FAB subtype M0 or M2) and demonstrate expression of the stem cell markers CD34 and CD117.^5,38^ Waraky et al. used retroviral transduction of *MNX1*-expressing constructs into murine fetal HSPCs and were able to induce AML.^39^ A possible link to leukemogenesis was described with the observation that *MNX1* activation resulted in reduced H3K4me1/2/3 and H3K27me3 levels providing increased chromatin accessibility.^39^

Our study challenges the dogma in AML that all reciprocal translocations lead to oncofusion proteins as an overestimated molecular mechanism in AML for gene activation. Future studies unraveling the molecular defects of t(7;12)(q36;p13) AML should focus on the targets of homeobox transcription factor *MNX1* rather than the oncofusion, as already initiated in some reports.^28,37,40^ However, also the role of genes on chromosome 19 (e.g. *DNA methyltransferase 1, DNMT1*, or *RNA polymerase II transcriptional elongation factor, ELL*), and effects of *ETV6* haploinsufficiency should be considered as co-factors in the leukemogenic process. ETV6 is a strong transcriptional repressor, and haploinsufficiency could result in reactivation of its target genes.^41^

## Data availability

All cell line sequencing data generated here are based on human reference GRCh37/hg19 and were deposited at the NCBI Gene Expression Omnibus (GEO).

## Acknowledgments

The authors would like to thank the Genomics and Proteomics Core Facility, the Omics IT and Data Management Core Facility of the DKFZ Heidelberg and the Center for Advanced Light Microscopy (CALM) at the Ludwig-Maximilians-University Munich for their excellent support. The work was in part supported by funds from the Helmholtz Foundation (ENHANCE Program) and German Research Foundation, SFB1074 subproject B11N (to CP and AR), Carreras Foundation (CP, ES) and FOR2674 subprojects A1, A6, A9 (CP, DBL), the Swedish Cancer Society (20 0925 PjF, CAN2017/461), the Swedish Childhood Cancer Foundation (PR2021-0025 and TJ2022-0017) and Västra Götalandsregionen (ALFGBG-431881), to LP and the SFB1064 subproject A17 (HL). Further support comes from the Helmholtz International Graduate School (AR, ES).

