## Supplemental File for "Altered enhancer-promoter interaction leads to *MNX1* expression in pediatric acute myeloid leukemia with t(7;12)(q36;p13)"

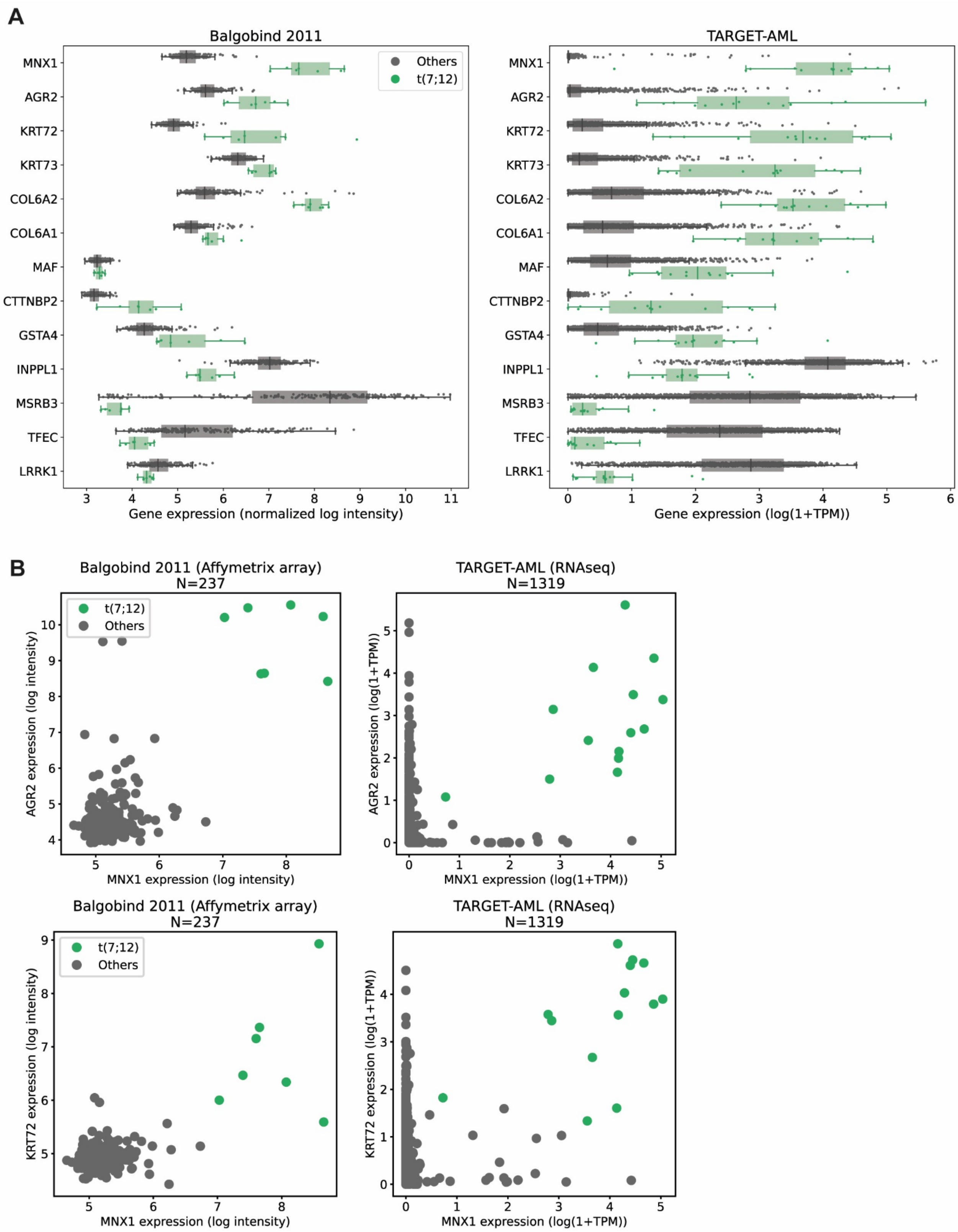


**Supplemental Figure 1: Differentially expressed genes in AML with or without t(7;12)(q36;p13)**

**A**, Boxplots for the expression levels (normalized log intensity or log(1+TPM) of the most differentially expressed genes between t(7;12) AML and all other samples, in the Balgobind and TARGET-AML cohorts. **B**, Scatter plot of the joint expression of *MNX1* and *AGR2*, or of *MNX1* and *KRT72*.

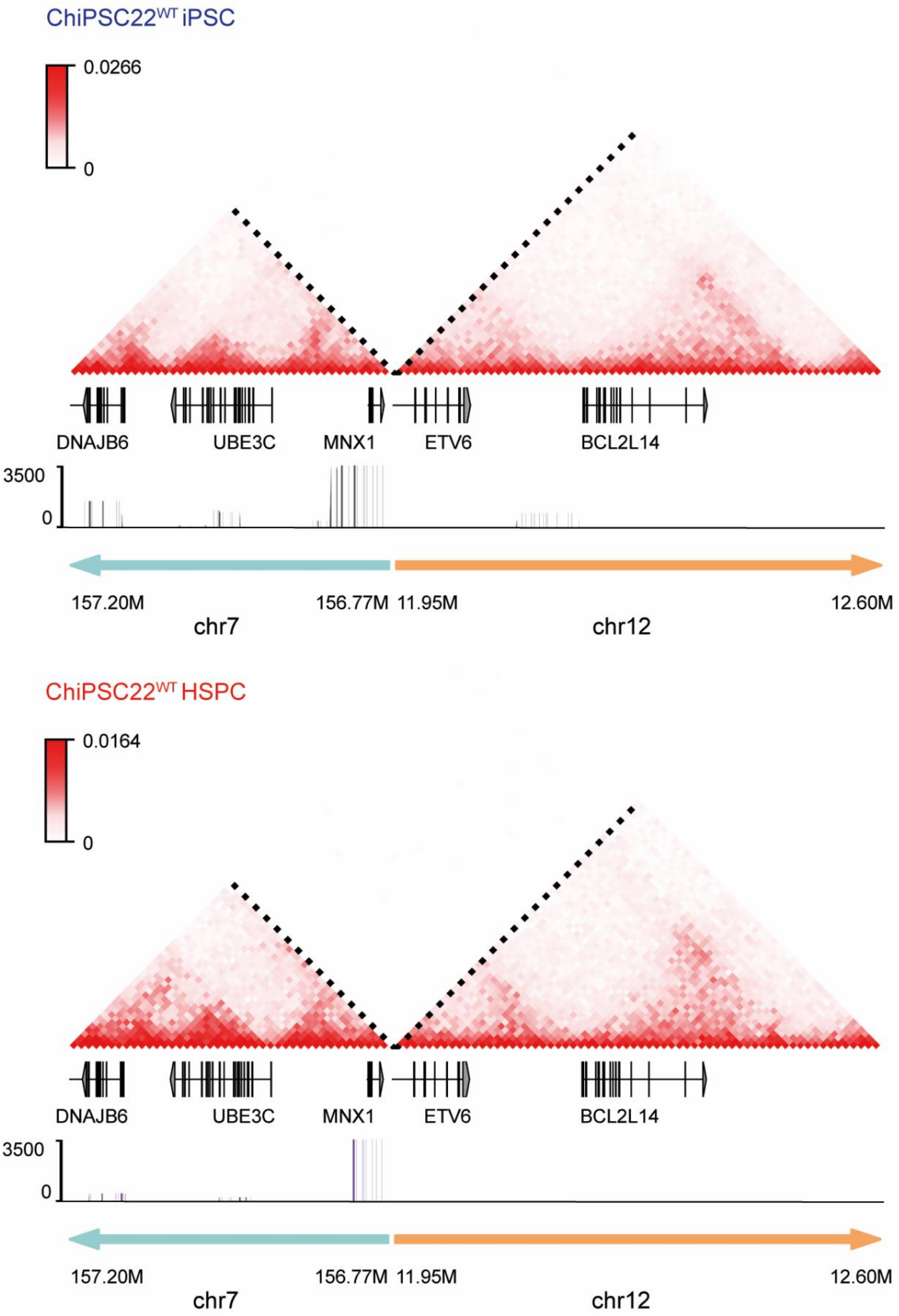


**Supplemental Figure 2: Interactions between *ETV6* region and *MNX1* promoter**
Chromatin interactions analyzed by Hi-C seq in the genomic region flanking the translocation breakpoint in the ChiPSC22$^{WT}$, either as iPSCs (top) or HSPCs (bottom). The tracks below show 4C-data with a viewpoint about 3 kb distal to *MNX1*. The range 0-3500 corresponds to read numbers in bins..

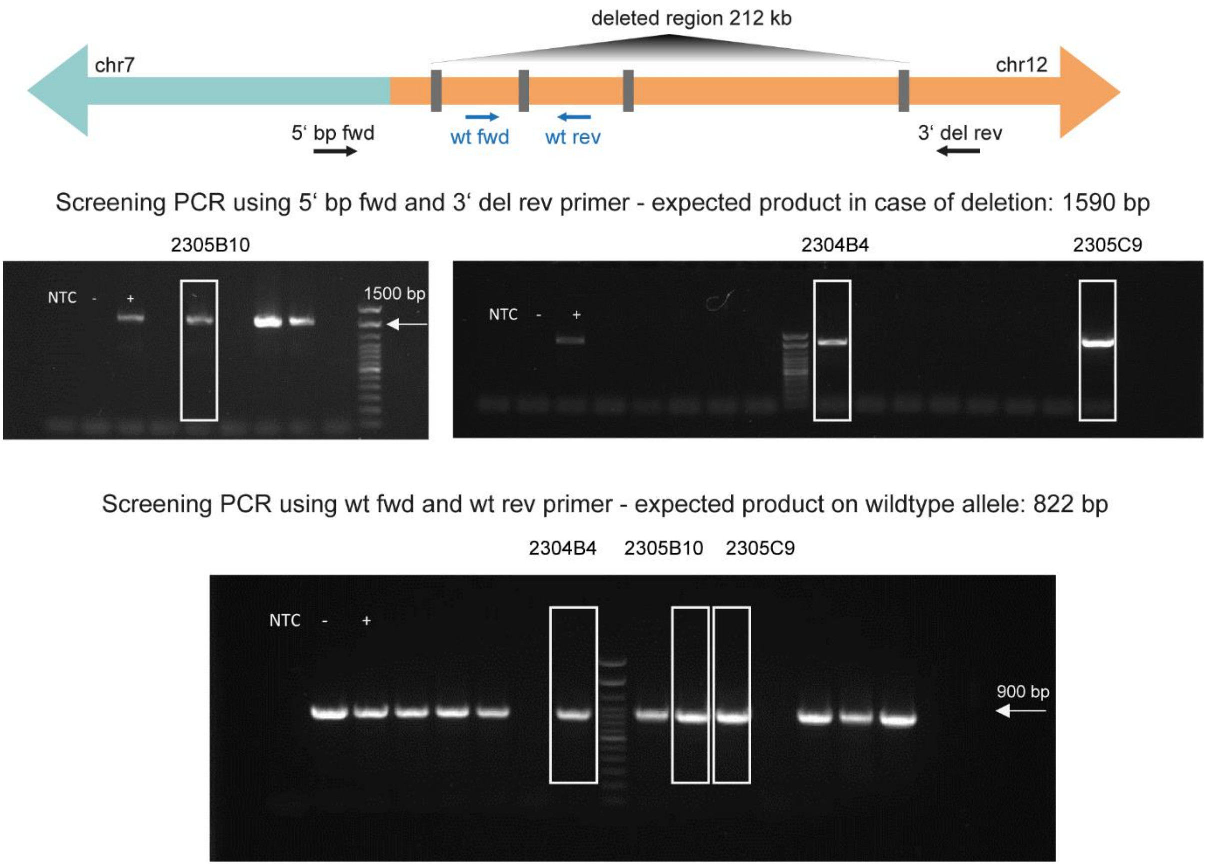


**Supplemental Figure 3: Validation of enhancer deletion experiments**

Scheme showing the location of the screening primers used to verify the presence of the deletion on the translocated allele and the presence of a wildtype *ETV6* region. Upper PCR results in 1590 bp product if enhancer deletion is present on the translocated allele and lower PCR results in 822 bp product if wildtype *ETV6* locus is present. Cell lines used in the study are highlighted in white box and showed PCR products in both PCRs.

### Supplemental Table 1: Pediatric AML expressing *MNX1*

| Sample ID | Cohort | Available cytogenetic information | age at diagnosis [month] |
|---|---|---|---|
| **T1** | Germany | 47,XY,**t(7;12)(q36;p13)**,+19[17]/47,idem,t(1;15)(p34;q21)[3]/48,idem,+8[2]. ish der(7)t(7;12)(5'TEL+),der(12)t(7;12)(5'TEL-)[19/19]. nuc ish 12p13(TELx2)(5'TEL sep 3'TELx1)[93/100] | 4 months |
| **T2** | Germany | 47,XY,**t(7;12)(q36;p13)**,+19[20] | 15 months |
| **T3** | Germany | 47,XX,**t(7;12)(q36;p13)**,+19[14]/46,XX[1] | 7 months |
| **T4*** | Sweden | 47,XY,**t(7;12)(q36;p13)**,+19[13]/46,XY[2] | 6 months |
| **TARGET-20-PARUNX** | TARGET-AML | 48,XX,+der(6)t(1;6)(q21;q27),**t(7;12)(q36;p13**),+19[18] [t(7;12) nuc ish ETV6 sep] | 77 months |
| **TARGET-20-PASIBG** | TARGET-AML | 47,XY**,t(7;12)(q36;p13)**,+19[20] | 25 months |
| **TARGET-20-PAWUTL** | TARGET-AML | 49,XX,+6,del(7)(q22),add(12)(p13.3),+19,+22[19]/46,XX[1] | 10 months |
| **TARGET-20-PAVXZL** | TARGET-AML | 47,XY,**t(7;12)(q36;p13)**,+19[17]/47,idem,dup(2)(q13q33)[6]/47,idem,del(11)(q14q25)[2] | 11 months |
| **TARGET-20-PAXMPG** | TARGET-AML | 47,XX,?add(7)(q32), cryp ins(7;12)(q36;p13.2p13.2),+19[20] | 11 months |
| **TARGET-20-PAXHGR** | TARGET-AML | 47,XX,**t(7;12)(q36;p13)**,+19[20] | 5 months |
| **TARGET-20-PAUYCA** | TARGET-AML | 46,XY[20] | 176 months |
| **TARGET-20-PAWNHH** | TARGET-AML | 47,XX,**t(7;12)(q36;p13)**,+19[20] | 17 months |
| **TARGET-20-PAVCJB** | TARGET-AML | 51,XY,+6,t(7;12;14)(q36;p13;q32.1),+8,+15,+19,+21[20] | 8 months |
| **TARGET-20-PAXEWS** | TARGET-AML | 47,XY**,t(7;12)(q36;p13)**,+19[20] | 6 months |
| **TARGET-20-PAVXPB** | TARGET-AML | 49,XY,+6,**t(7;12)(q36;p13)**,+16,+19[20] | 24 months |
| **TARGET-20-PAWBTJ** | TARGET-AML | 46,XX,der(7)del(7)(q22q36)**t(7;12)(q36;p13.3)**,der(12)t(7;12)(q36;p13.3)[20] | 9 months |
| **TARGET-20-PAVFDW** | TARGET-AML | 46,XX[20] | 209 months |
| **TARGET-20-PAVAAM** | TARGET-AML | 46,XX,t(6;11)(q27;q23)[20] | 56 months |
| **TARGET-20-PAWNYK** | TARGET-AML | 46,XX,**t(7;12)(q36;p13)**[7]/46,XX[13] | 20 months |
| **TARGET-20-PAVNGY** | TARGET-AML | 46,XY[20] | 161 months |
| **TARGET-20-PAWYKA** | TARGET-AML | 46,XY[30] | 102 months |
| **TARGET-20-PAVDGM** | TARGET-AML | 46,XY[29] | 199 months |

| | | | |
|---|---|---|---|
| **TARGET-20-PAWMLK** | TARGET-AML | 46,XX[60] | 194 months |
| **TARGET-20-PAUYZY** | TARGET-AML | 46,XY,del(11)(q21q24),der(11)ins(11;11)(p15;q21q23)[11]/46,XX[9] | 15 months |
| **TARGET-21-PASVJS** | TARGET-AML | 46,XX,[20] | 63 months |
| **TARGET-20-PAUSBP** | TARGET-AML | 46,XY,t(6;11)(q27;q23)[18]/46,XY[2] | 140 months |
| **TARGET-20-PAXKAL** | TARGET-AML | 46,XY,t(6;17)(q21;p13),der(7)t(7;7)(q36;p13),add(9)(p13),add(18)(q23)[cp3]/46,sl,del(1)(q32)[2]/46,sl,der(7)t(?4;7)(q21;q?32)[3]/46,sdl2,der(7)del(7)(p13)t(7;7)(q36;p13),add(19)(q13.3)[9]/90,sdl3x2,-X,-21[cp2] | 176 months |
| **gsm446011**** | Balgobind 2011 | **t(7;12)** | <18 months |
| **gsm445924**** | Balgobind 2011 | **t(7;12)** | <18 months |
| **gsm446153**** | Balgobind 2011 | **t(7;12)** | <18 months |
| **gsm445951**** | Balgobind 2011 | **t(7;12)** | <18 months |
| **gsm445920**** | Balgobind 2011 | **t(7;12)** | <18 months |
| **gsm446155**** | Balgobind 2011 | **t(7;12)** | <18 months |
| **gsm446154**** | Balgobind 2011 | **t(7;12)** | <18 months |

*qRT-PCR resulting in 4-fold higher *MNX1* than *ABL1* expression with assays Hs00907365 (*MNX1*, $C_t$ =24.4) and Hs01104728 (*ABL1*, $C_t$=26.4, reference).
**Microarray expression analysis
All other samples analyzed by RNAseq, and the first six samples analyzed by WGS.

**Supplemental Table 2: Known AML driver genes**

*ASXL1*
*ASXL2*
*BCOR*
*CEBPA*
*CEBPG*
*CREBBP*
*DNMT3A*
*DNMT3B*
*ETV6*
*EZH2*
*FLT3*
*GATA2*
*IDH1*
*IDH2*
*JAK2*
*JARID2*
*KAT6A*
*KDM3B*
*KDM6A*
*KIT*
*KMT2A*
*KMT2C*
*KMT2D*
*KMT2E*
*KRAS*
*MED12*
*NCOR1*
*NCOR2*
*NF1*
*NOTCH1*
*NOTCH2*
*NPM1*
*NRAS*
*NSD1*
*PHF6*
*PTPN11*
*RB1*
*RUNX1*
*SF3B1*
*SMARCA2*
*SMARCA4*
*SMC1A*
*SMC3*
*SRSF2*
*STAG2*
*SUZ12*
*TET1*
*TET2*
*TP53*
*U2AF1*
*WT1*
*ZRSR2*

**Supplemental Table 3: TaqMan gene expression assays (Applied Biosystems)**

| **Gene name** | **Assay ID** |
|---|---|
| *MNX1* | Hs00907365_m1 |
| *AGR2* | Hs00356521_m1 |
| *MMP9* | Hs00957562_m1 |
| *CRISP3* | Hs00195988_m1 |
| *MAF* | Hs04185012_s1 |
| *GUSB* | Hs99999908_m1 |

**Supplemental Table 4: Antibodies**

| Antigen | Source | Purpose | Immunogen | Manufacturer and product number | RRID | Amount or dilution used |
|---|---|---|---|---|---|---|
| β-actin | Mouse monoclonal | Western | β-actin | Santa Cruz sc-47778 HRP | AB_2714189 | 1:10000 |
| MNX1 | Rabbit polyclonal | Western | aa 6-43 | Thermo Fisher PA-67195 | AB_2662922 | 1:500 |
| Rabbit immunoglobulins | Goat | Western | HRP-linked anti rabbit IgG | Cell Signaling 7074 | AB_2099233 | 1:5000 |
| Mouse immunoglobulins | Goat | Western | HRP-linked anti mouse IgG | Cell Signaling 7076 | AB_330924 | 1:5000 |
| H3K4me1 | Rabbit polyclonal | ACT-seq | aa 1-100 (mono methyl K4) | Abcam ab8895 | AB_306847 | 0.8 µg |
| H3K27ac | Rabbit polyclonal | ACT-seq | aa 1-100 (acetyl K27) | Abcam ab4729 | AB_2118291 | 0.8 µg |
| H2B | Rabbit monoclonal | ACT-seq | Synthesized peptide from yeast H2B | Boster Biological Technology M30930 | AB_2924769 | 0.8 µg |
| IgG | Rabbit serum | ACT-seq | not applicable | Millipore PP64B | AB_97852 | 0.8 µg |

### Supplemental Table 5: Primers used for 4C

| Viewpoint region | Viewpoint primer | Sequence[1] | hg19 primer coordinate | Touchdown PCR | PCR cycles |
|---|---|---|---|---|---|
| *MNX1* 3 kb upstream | HindIII_4cMNX1_R1 | GTCTCGTGGGCTCGGAGATGTGTATAAGAGACAGGGCTAGGTGTACCTGAAAGCTT | chr7:156805780-156805801 | | |
| | DpnII_4cMNX1_F1 | TCGTCGGCAGCGTCAGATGTGTATAAGAGACAGGACCAAATGGACTCTAAGGAGC | chr7:156806553-156806574 | 63°C-58°C | 35 |

[1]Genome-specific sequence corresponding to hg19 coordinate is underlined; break point location is chr7:156778230.

**Supplemental Table 6: Two-color FISH oligonucleotides**

See separate file for Supplemental Table 6

**Supplemental Table 7: PCR primers to confirm engineered deletion on chr12**

| Name | Sequence 5' to 3' | hg19 position |
|---|---|---|
| 5' bp fwd | CTTCACTCTATGCCTCCCTACCTGG | chr7: 156778089-156778113 |
| 3' del rev | GCAAGTGCCTATGCAAACCA | chr12: 12164774-12164793 |
| wt fwd | CCAGGTGCAAGGCTCTATCC | chr12: 11989035-11989054 |
| wt rev | TTGAACAAGAATGGCGCTGC | chr12: 11989909-11989928 |
| 5' del crRNA | TCAGGCACAATCCTTTGTACTGG | chr12: 11951006-11951028 |
| 3' del crRNA | AGCACATCCCAACTGGCACTGGG | chr12: 12164572-12164594 |
| ssODN | AAGGATAAGATTTTGAGATTATCTTTCCAAGGAAAACTTATCAACTATCTATTATGTATCAGGCACAATCCTTTGGCCAGTTGGGATGTGCTGTGCACACCAAGTGCCCCTAGCTCCTCACCTACCTCTGGGATCACTGACAGTCCTGCT | black: chr12: 11950948-11951022; red: chr12: 12164578-12164652 |

**Supplemental Table 8: Differentially expressed genes in t(7;12)(q36;p13) AML cases from the Balgobind [40] and the TARGET [37] cohorts**

See separate file for Supplemental Table 8
